# “Sonohistology”: Ultrasonographic Tissue Characterization in the Diagnosis of Hepatic Steatosis

**DOI:** 10.1101/2022.07.24.501305

**Authors:** Luís Jesuino de Oliveira Andrade, Luís Matos de Oliveira, Maria Helena Ferreira Andrade, Gabriela Correia Matos de Oliveira

## Abstract

**Objective:** To correlate ultrasound imaging and histology in hepatic steatosis.

**Material and method:** Liver biopsy slides of cases in which hepatic steatosis were classified as mild, moderate and severe was evaluated, determining the number of fat cells per microscopic field from the histological sections. The number of fat cells per field was correlated with the degree of hepatic steatosis suggested on ultrasound examination.

**Results:** Mean number of fat cells per microscopic field was 50 in mild steatosis, 85 in moderate steatosis, and greater than 150 fat cells in severe steatosis. A significant correlation was observed between echogenicity and the number of fat cells, i.e. the more fat cell the higher the proportion of sound waves incident perpendicularly, being reflected back to the transducer (hyperechogenicity), while in normal liver tissue there is a greater dispersion of the sound wave (hypoechogenicity).

**Conclusion:** The main interface of echo reflection in steatotic liver tissue is the boundary between the normal hepatocyte and fatty infiltration.

## Introdução

Hepatic fatty infiltration is defined when more than 5% of hepatocytes have large fat vacuoles, and it is estimated that nonalcoholic fatty infiltration of the liver affects about 1 billion people worldwide, comprising about 19-46% of liver diseases occurring in the Western world.^1,2^

Liver biopsy, despite possible harms, is considered to be the current gold standard in diagnosing hepatic steatosis.^3^ Hepatic fatty infiltration is classified histologically according to the percentage of large or medium fat droplets in the hepatocyte: less than 5% normal, between 5-33% mild steatosis (Grade 1), between 34-66% moderate steatosis (Grade 2), and greater than 66% severe steatosis (Grade 3).^4,5^

Imaging methods are commonly used for evaluation of hepatic steatosis and ultrasonography is the initial imaging method used in the identification and classification of hepatic fatty infiltration, because it allows subjective assessment of the degree of fatty infiltration in the liver.^6^ Studies have demonstrated that ultrasonography presents quite variable sensitivity and specificity, which limits this method in the diagnosis of this disease.^7^ However, a meta-analysis, has demonstrated that ultrasonography is an accurate and reliable imaging method in the identification of moderate and severe hepatic steatosis when compared with histology.^8^ Hepatic fatty infiltration is classified sonographically into: Normal (Grade 0) when the hepatic echographic texture is normal; Mild (Grade 1) when a level, diffuse, non-attenuating increase in hepatic echogenicity is observed, with normal visualization of the intrahepatic vessels and diaphragm; Moderate (Grade 2) when a moderate increase is observed with attenuation of the acoustic beam, with impaired visualization of the diaphragm and hepatic vascularization; Severe (Grade 3) when a severe and diffuse increase in hepatic echogenicity is observed, without visualization of the intrahepatic vessels and posterior liver contours.^9^

Therefore, ultrasonography in assessing the degree of hepatic fatty infiltration would be a useful noninvasive method in predicting liver histology, especially in cases where biopsy is contraindicated. The aim of this study was to compare the degree of echogenicity of hepatic fatty infiltration on two-dimensional ultrasonography with the histological degree of hepatic fatty infiltration based on the number of fat cells per histological field.

## Methods

Photographs of liver biopsy slides with normal liver, mild, moderate and severe steatosis were obtained in a specialized pathological anatomy laboratory, evaluating the number of fat cells per microscopic field from the histological sections. The number of fat cells per field was correlated with the degree of hepatic steatosis suggested on ultrasound examination.

Estimation of the degree of hepatic fat infiltration by two-dimensional ultrasonography was obtained using ultrasound features that included the brightness, contrast between the liver and the adjacent kidney, acoustic beam attenuation, the visibilization of intrahepatic vessels, and the posterior contours of the hepatic lobe.

According to the Resolution CNS 510/2016, the study did not require the submission of the ethics committee, due to the fact that the research aimed to deepen the theoretical understanding of pathologies of clinical practice

## Results

A significant correlation occurred between echogenicity and the number of fat cells. The greater the number of fat cells the greater the proportion of sound waves incident perpendicularly, being reflected back to the transducer (hyperechogenicity).

Using the basic physical principles of ultrasonography ^10^ for the analysis of low and high echogenicity we present a combination of parameters (Table 1). Sound reflection results from the difference in acoustic impedance at the boundary between two media, while sonic energy reduction results from sound absorption and sound dispersion.

**Table 1.** Physical parameters of echogenicity

|  | Sound absorption | Scattering | Difference of acoustic impedance at interfaces |
| --- | --- | --- | --- |
| Low echogenicity | High | High | Low |
| High echogenicity | Low | Low | High |

A significant correlation was observed between echogenicity and the number of fat cells, i.e. the more fat cell the higher the proportion of sound waves incident perpendicularly, being reflected back to the transducer (hyperechogenicity), while in normal liver tissue there is a greater dispersion of the sound wave (hypoechogenicity).

Ultrasonography and histology of a normal liver: ultrasound longitudinal section demonstrating liver tissue of normal echogenicity (Figure 1).

**Figure 1.**
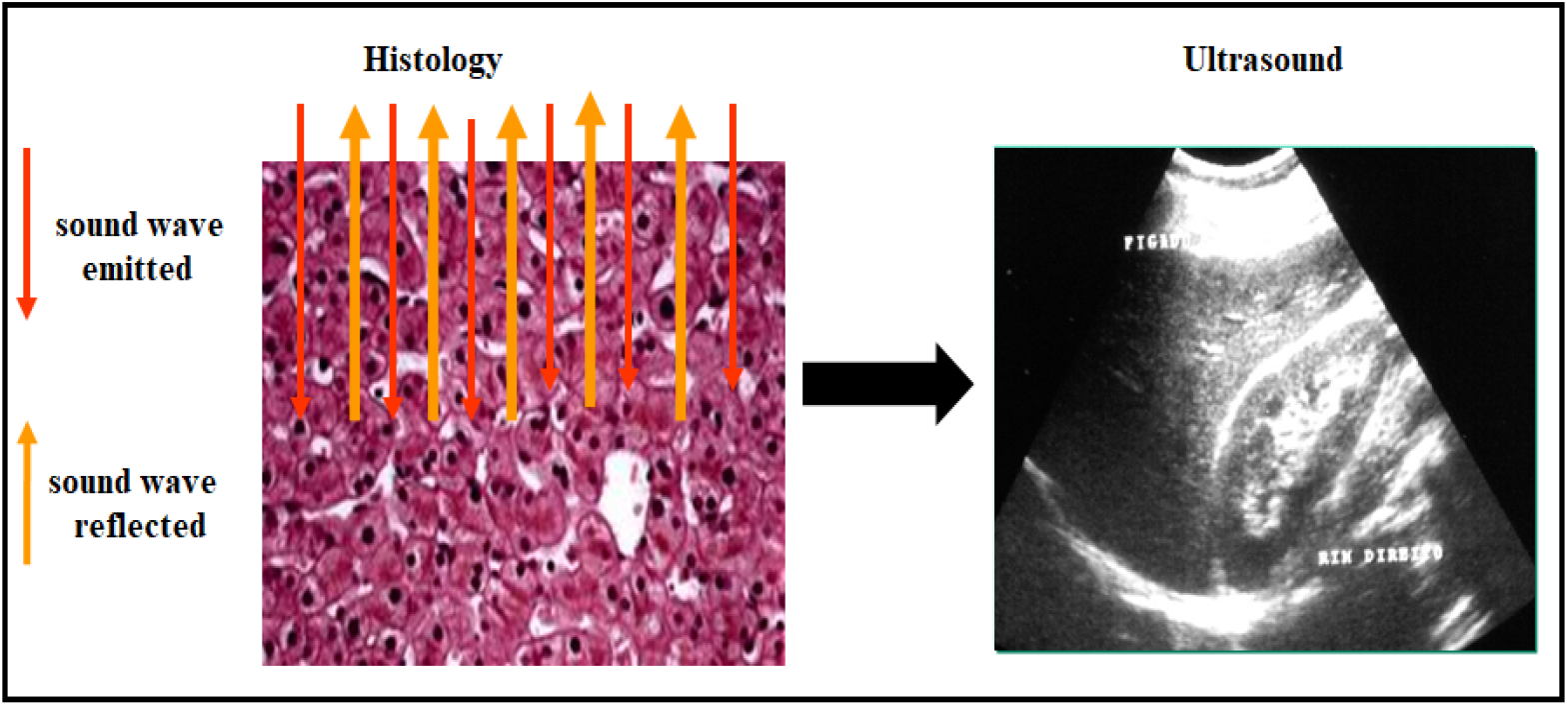
Ultrasonography and histology of a normal liver. **Source**: search result

In the histological evaluation of the mild hepatic fatty infiltration, the mean number of fat cells per microscopic field was 50 cells (Figure 2).

**Figure 2.**
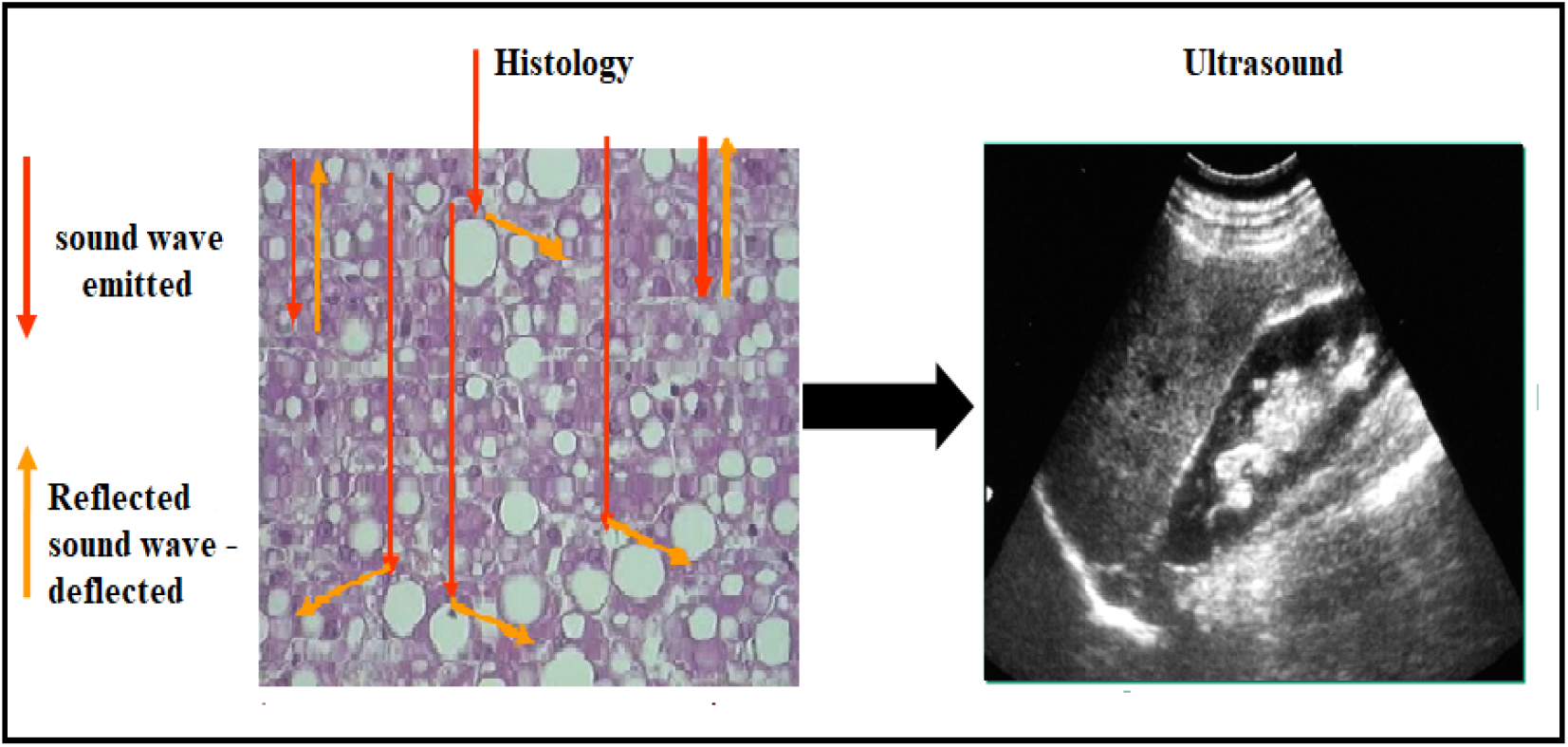
Ultrasonography and histology of the mild hepatic fatty infiltration. **Source**: search result

In the histological evaluation of the moderate hepatic fatty infiltration, the mean number of fat cells per microscopic field was 85 cells (Figure 3).

**Figure 3.**
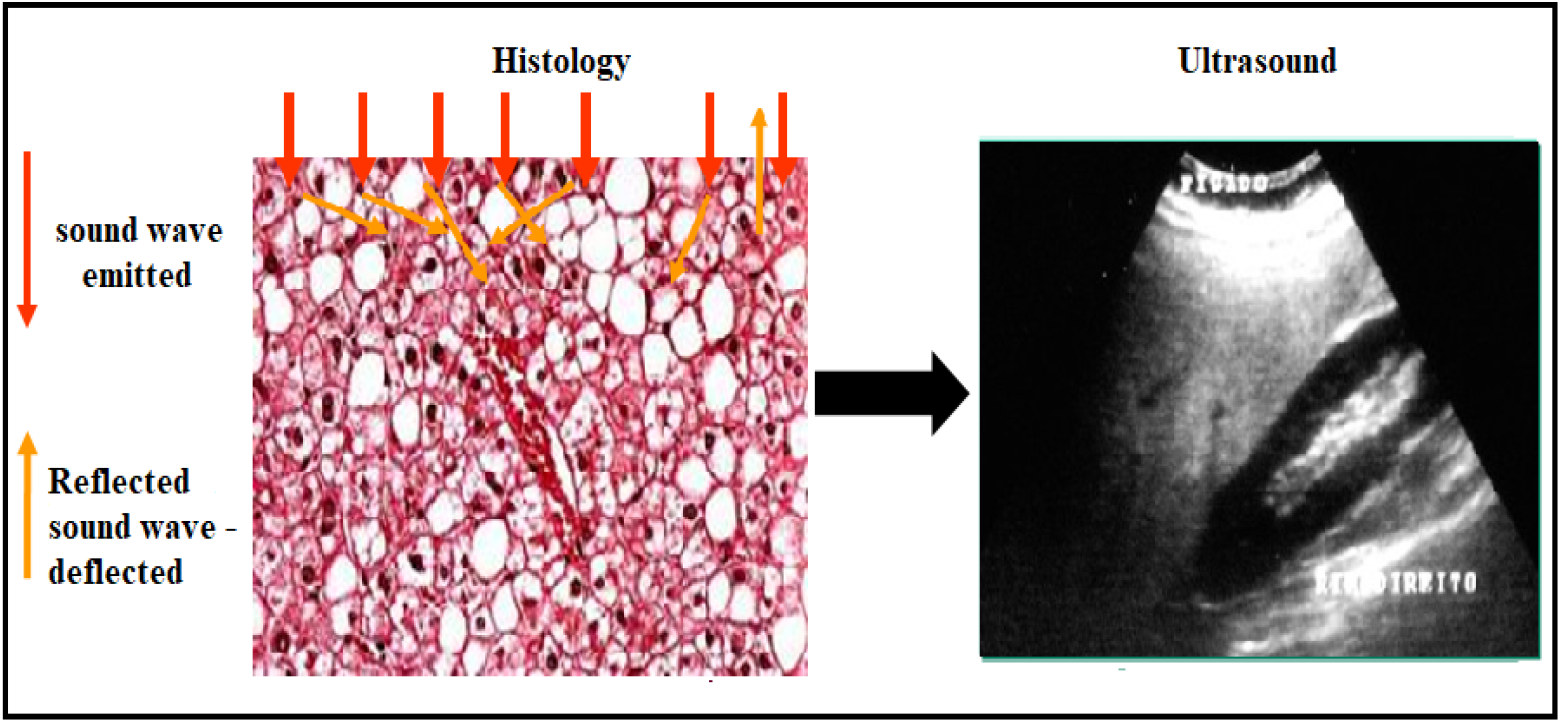
Ultrasonography and histology of the moderate hepatic fatty infiltration. **Source**: search result

In the histological evaluation of the severe hepatic fatty infiltration, the mean number of fat cells per microscopic field was > 150 cells (Figure 4).

**Figure 4.**
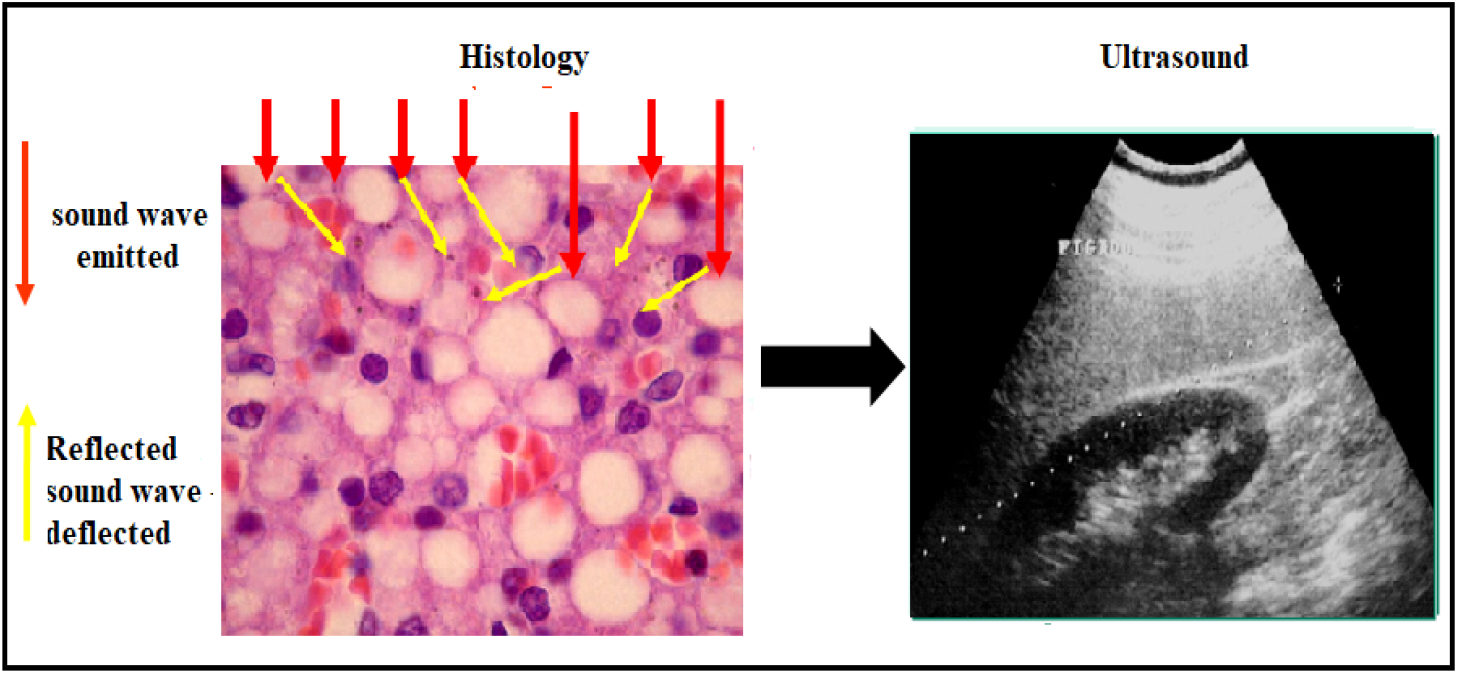
Ultrasonography and histology of the severe hepatic fatty infiltration. **Source**: search result

## Discussion

Our results demonstrate that the degree of hepatic echogenicity on ultrasound shows a quantitative correlation of histologic severity in hepatic fatty infiltration based on the number of fat cells presenting per fields on histologic sections, “sonohistologically” characterizing the nonalcoholic fatty liver disease.

“Sonohistology” in sonographic tissue characterization in the diagnosis of hepatic fatty infiltration can be defined as the correlation between the subjective quantitative parameters of hepatic echogenicity and the objective histological parameters of the liver tissue analyzed by pathological examination.

Several imaging studies have been performed in order to characterize the tissue of hepatic fatty infiltration, such as dual-energy computed tomography and magnetic resonance imaging using neural networks, in addition to ultrasound.^11-13^ We use the physical principles of ultrasound to analyze normal tissue that is characterized histologically by higher cellularity translated on ultrasound by an average echogenicity slightly more hypoechogenic than the spleen and brighter than the adjacent kidney. Thus, in normal liver tissue there is a greater dispersion of the sound wave (hypoechogenicity).

Non-alcoholic fatty liver disease is the most common pathology found in the liver today. Carefully applied ultrasonography has been found to be a good diagnostic method. Liver tissue with fat infiltration is characterized by increased echogenicity, compared to the echogenicity of the adjacent spleen and kidney, due to the percentage of fat cells present in this tissue. With this occurs an attenuation of the ultrasound beam with reduced sound intensity through the liver tissue changing the absorption and dispersion of sound with consequent divergence of the acoustic beam. The attenuation of sound in the insonated structure reduces the visualization of details of the liver structures.^14^ Hepatic fatty infiltration is categorized based on the percentage of fat within the hepatocytes: grade 0 (healthy, <5%), grade 1 (mild, 5%-33%), grade 2 (moderate, 34%-66%), and grade 3 (severe, >66%).^15^ Our study observed that the greater the number of fat cells the greater the proportion of sound waves that incur perpendicularly and are reflected back to the transducer (hyperecogencity). Thus, a 5%-33% percentage of fat cells in the hepatocyte corresponded on average to 50 adipocytes per field, between 34%-66% corresponded on average to 85 adipocytes per field, and more than 66% of fat cells in the hepatocyte corresponded to more than 150 adipocytes per field in the histological slides evaluated.

Based on this study we can consider the echogenicity of the liver tissue as a measure of its cellular content. Normal echogenicity demonstrates the existence of normal hepatocytes with normal homogeneous echographic pattern thus ruling out other pathologies such as fatty infiltration. Thus, the main echo reflection interface in steatotic liver tissue is the boundary between normal hepatocyte and fatty infiltration.

